# Impact of long-term storage on mid-infrared spectral patterns of serum and synovial fluid samples of dogs with osteoarthritis

**DOI:** 10.1101/2022.10.02.510573

**Authors:** Sarah Malek, Federico Marini, J T. McClure

## Abstract

**Objective:** To evaluate impact of long-term storage on mid-infrared (MIR) spectral patterns of serum and synovial fluid (SF) of dogs with knee OA and controls.

**Design:** Serum (52 OA and 49 control) and SF (51 OA and 51 control) samples from dogs that had been in short-term (<3 years) frozen state (−80°C) had their MIR spectra obtained. The remaining aliquots were maintained in long-term (>5 years) frozen state before having MIR spectra acquired under the same testing conditions. Multi-level simultaneous component analysis was used to evaluate the effect of time. Partial least squares discriminant analysis was used to compare performance of predictive models built for discriminating OA from control spectra from each time point.

**Results:** Median interval of storage between sample measurements was 5.7 years. Spectra obtained at two time points were significantly different (*P* <0.0001), however, contribution of sample aging accounted for only 1.61% and 2.98% of serum and SF profiles’ variability, respectively. Predictive models for discriminating serum of OA from controls for short-term storage showed 87.3±3.7% sensitivity, 88.9±2.4% specificity and 88.1±2.3% accuracy, while, for long-term storage, values of the same figures of merit were 92.5±2.6%, 97.1±1.7% and 94.8±1.4%, respectively. Predictive models based on short-term stored SF spectra had 97.3±1.6% sensitivity, 89.4±2.6% specificity and 93.4±1.6% accuracy, while the values for long-term storage 95.7±2.1%, 95.7±0.8% and 95.8±1.1%, respectively.

**Conclusions:** Long-term storage of serum and SF results in significant differences in spectral variables, however, these changes do not significantly alter the performance of predictive algorithms for discriminating OA samples from controls.

## 1. Introduction

Biobanking biological fluid samples (e.g., blood, urine, joint fluid) is an essential component of prospective clinical studies and a means of maximizing efficiency of using obtained samples. These biological samples and the contents therein (i.e., cellular, molecular) are at risk of being compromised during the pre-analytical phase of projects. The pre-analytical phase is where sample collection technique, handling and storage protocols may inadvertently alter the components of interest. Cryopreservation of biological fluids requires standardized protocols, reliable inventory, and specialized facilities to preserve the valuable samples. Factors that have been shown to impact stability of the samples during the storage phase are temperature, storage time and freeze-thaw cycles. The impact of these factors on various biomarkers and sample components have been investigated in previous studies(1–3). However, recommendations for optimal storage variables vary depending on stability of the target components (e.g., molecule being measured) and measurement technique. Use of mid-infrared (MIR) spectral patterns of biological fluids to detect variability between disease states as a novel approach to disease pattern recognition has been investigated with an increasing rate in the past few decades(4–12). The spectral pattern of a given sample is the sum of all MIR light absorbance by the infrared-active molecular bonds within the sample that is displayed as a unique waveform (spectrum)(13). Fourier-transform infrared spectroscopy (FTIR) is one of the techniques utilized in acquiring MIR spectra of biological samples that is adjuvant free, simple and requires small volume of samples(5–9). Use of the spectral variables based on this method has been successfully used in differentiating various types of arthritis in humans(4). Research in experimental and clinical models of osteoarthritis (OA) in various joints of animal models has also been able to demonstrate ability to distinguish OA from control samples with high accuracy rates(5–9). Shortterm (< one year) longitudinal studies using spectral variables in serum and joint fluid of animals have shown reproducible spectral characteristics of OA that despite changes overtime, remain distinguishable from control samples(7–9). Studies investigating long-term changes in clinical OA in larger animal models (e.g., dogs) and human models require at least several years that warrants biobanking of collected biological samples to allow for batch analysis. These biobanked biological samples (e.g., blood, synovial fluid0 are typically stored in −80 degrees Celsius for long-term storage to minimize the impact of time. However, the impact of long-term storage on the spectral pattern of these biological samples has not been previously investigated. The first aim of this study was to evaluate whether long-term storage results in changes in the MIR spectral pattern of serum and synovial samples of dogs with and without knee OA as measured by FTIR spectroscopy. The second aim of this study was to evaluate the impact of any observed changes due to long-term storage on the ability to discriminate between the serum and synovial fluid spectra patterns of dogs with and without knee OA. The hypothesis of the study was that long-term storage will have minimal impact on MIR spectral pattern of serum and synovial fluid of dogs with or without knee OA.

## 2. Methods

### 2.1. Samples

The animal Care Committee of the University of Prince Edward Island (#11-062) had approved enrollment of dogs for the original sample collection with informed consent signed by respective owners for an earlier study (8, 9). The sample size used in the current study was based on available number of samples from the original project that had two arms investigating serum and synovial fluid samples from client-owned dogs with OA secondary to naturally occurring degenerative (non-traumatic) cranial cruciate ligament tear in one or both knees and controls(8, 9). The controls were otherwise healthy, adult dogs with no orthopedic abnormalities with both knees free of gross abnormalities upon evaluation immediately after euthanasia for reasons unrelated to the study. Briefly, venous blood samples had been collected and serum separated and saved in 0.5-1ml aliquots in cryovials (Nalgene Cryogenic tubes, VWR International, Batavia, IL, USA) and preserved at −80°C until batch analysis. Synovial fluid samples had been collected for the original study using aseptic technique from knees of dogs with cranial cruciate ligament tear in the OA group under general anesthesia immediately prior to surgical intervention to treat knee instability (i.e., tibial plateau leveling osteotomy and knee joint inspection via arthrotomy or arthroscopy), and from healthy knees in the control group immediately after euthanasia. The synovial samples were also saved in 0.5 ml aliquots in cryovials and preserved at −80°C until batch analysis. After the completion of the original project, the unused aliquots of serum and synovial samples were maintained in −80°C storage for a minimum of 5 years.

The available sample inventory was reviewed for OA and Control samples with adequate volume left for analysis. For the OA group the serum and synovial fluid samples were those obtained prior to any surgical intervention. If more than one aliquoted sample was available, the clearest was selected (i.e., non-to minimal blood contaminated for synovial fluid and none to minimal hemolysis for serum samples). Only one serum sample per dog was selected. In the OA group, even if both knees were affected, only one sample from one knee per dog was included in the study. In the control group, both healthy knees of each dog had been sampled, but preferentially only one sample from each dog was included.

The age of samples at the time of first spectral analysis was calculated based on the date sample was obtained from the dog and the time when the sample was thawed, and the first spectral analysis was performed. Samples from this initial storage period are referred to as “short-term storage” samples. The age of samples at the time of second spectral analysis after being in storage for a minimum of 5 years was calculated based on date of sample acquisition from the dog and the time when the sample was run for the second time. Samples from the longer storage period are referred to as “long-term storage samples”. The interval between measurements for the short-term and long-term storage was calculated based on the dates the MIR spectra of each sample was obtained.

### 2.2. FTIR spectroscopy

At the time of first spectral acquisition (i.e., after short-term storage), serum and synovial fluid samples were thawed at 22°C and dried films were prepared as described previously (8, 9, 14). For each sample, an aliquot was drawn and diluted in a potassium thiocyanate (KSCN) (SigmaUltra, Sigma-Aldrich Inc, St Louis, MO) solution (4 g/L) at a 2:1 serum/SF-to-KSCN ratio (40:20 μL). KSCN was used as an internal control. For each sample, replicate (8 μL per replicate) dry films were made on a silicon microplate (5). After drying at room temperature (20-22 °C), the microplate was mounted on a multi-sampler (HTS-XT, Autosampler, Bruker Optics, Milton, ON, Canada) interfaced to the infrared spectrometer (Tensor 37, Bruker Optics). Mid-infrared absorbance spectra in the range of 400 to 4,000 cm^-1^ was recorded with proprietary software (OPUS software, version 6.5, Bruker Optics, GmbH, Ettlingen, Germany). For each acquisition, 512 interferograms (scans) were accumulated and Fourier transformed to generate a spectrum with a nominal resolution of 4 cm^-1^ (5, 6, 15). Six (short-term storage) or three (long-term storage) replicates (due to limited volume of available samples) were prepared and analyzed for each sample and the corresponding spectra were averaged prior to the successive data processing. At each measurement time point, all the spectra were acquired within a short time span (~ 10 days).

### 2.3. Statistical analysis

Analyses of non-spectral data were performed using SPSS software (IBM SPSS Statistics, v. 25). Variables without a normal distribution were described by their median and interquartile range (IQR). All acquired serum and synovial fluid spectra files from both time-points were imported in MATLAB^®^ (R2015b (8.6.0.267246); The Mathworks, Natick, MA) for the successive data processing which was carried out utilizing in-house written scripts. For each sample, at each time point, the average of the replicate spectra was used for analysis. Prior to any modeling, the data were preprocessed by first derivative (using Savitzky-Golay algorithm (16) with 19 points window and second order polynomial) followed by mean centering.

Modeling was conducted in two different stages. At first, multilevel simultaneous component analysis (MSCA) was used to verify whether there could be any significant difference between the spectra of the same samples measured after short- and long-term storage. Indeed, MSCA can be considered as a multivariate generalization of repeated measurements ANOVA (17, 18). In particular, the overall (preprocessed) spectral matrix ***X*** is partitioned into the individual contributions of between-sample (***X***_*b*_) and within-sample (***X***_*w*_) systematic sources of variability plus the residuals (***X***_*res*_, i.e., the variation not accounted for by the model), according to:

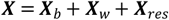

In particular, the effect of sample aging on the spectroscopic signal is related to the within-sample variation described by ***X***_*w*_ and can be quantified as the sum of squares of the elements in that matrix. Significance of the contribution is evaluated by comparing such effect with its distribution under the null hypothesis, which is estimated by means of a permutation test. On the other hand, if the effect is found to be significant, its impact on the spectroscopic signal can be interpreted by PCA of the associated matrix ***X***_*w*_. In the present study, MSCA analysis was conducted independently on the serum and synovial fluid samples.

In a second stage, to verify whether MIR analysis of synovial fluid or serum samples could provide the basis for a reliable discrimination between OA and control even after long-term storage, and if the same spectroscopic markers could be found as when analysis was carried out after short-term storage, supervised pattern recognition (classification) models were built and validated (19). Due to its ability to deal with highly collinear predictors (e.g., spectral variables), partial least squares discriminant analysis (PLS-DA) was selected (20, 21). The PLS-DA method is based on projecting the data onto a low-dimensional sub-space of latent variables, which are relevant to highlight differences between the groups. Accordingly, model building requires to identify the optimal dimensionality of such subspace. On the other hand, the quality of the predictive models, which can be summarized by different figures of merit, such as sensitivity, specificity, overall classification accuracy and the area under the ROC curve (AUC), needs to be evaluated on samples not used for model development, in order to avoid overoptimistic results. Accordingly, to build the models and validate the prediction results and the identified spectroscopic markers, a repeated double cross-validation (rDCV) procedure coupled with permutation tests was adopted (22, 23). In rDCV the available samples are split according to two nested loops of cross-validation: the outer loop samples are treated as external validation samples, which do not take part either in model building or in model selection; indeed, optimization of the model parameter is carried out on the inner loop samples. Furthermore, to rule out any possibility of obtaining good results just due to chance correlations, permutation tests were used to non-parametrically evaluate the null distributions of the classification figures of merit providing *P* values to estimate the significance of the observed discrimination (22). Statistical significance was set at *P* <0.05. In The analyses was conducted independently to report model performance based on whether serum or synovial fluid samples were used.

## 3. Results

There were 52 serum and 51 synovial fluid samples available from the 70 dogs originally in the OA group, while 49 serum and 52 synovial fluid samples were available from the 50 dogs in the control group. In the control group, one dog did not have adequate serum sample available. Two control dogs did not have adequate synovial fluid samples; therefore, the additional three samples were selected from the contralateral knees of three of control dogs that had adequate samples from both knees.

The median (IQR) of time in frozen state before the first run (short-term storage) and the second run (long-term storage) were 1.1 (1.1) and 6.8(1.1) years respectively. The time interval between the first and second measurement was 5.7 (0.03) years. At first, MSCA was used to evaluate whether there could be any significant difference in the spectral profiles of the samples after short or long-term storage. At first, serum samples were considered. When the overall variability in the preprocessed spectral profiles was partitioned according to the ANOVA scheme, it was found that only 1.61% of the total variability among the serum spectra was ascribable to the effect of aging (between-samples differences accounted for 77.23% of the total variance, while the remaining part is residual variation, associated to random error). The variability observed between the two time points was not impacted by the type of sample (OA versus control). Although the relatively low amount of spectral variance was associated with the differences between short-term and long-term storage, permutation testing indicated that these differences were statistically significant (*P* <0.0001). Principal component analysis (PCA) analysis of the effect matrix resulting from the ANOVA decomposition indicated that long-term stored serum samples are characterized, on average, by a lower intensity of most of the peaks. The same analysis was then conducted on the synovial fluid samples. Also in this case, when the overall variability in the preprocessed spectral profiles was partitioned according to the ANOVA scheme, it was found that the largest part of the spectral variance was due to between-samples differences (85.54%), with only 2.98% of the total variability among the spectra corresponding to the effect of aging. Permutation test indicated that the effect of sample ageing on the spectral signature, though relatively small, could still be deemed statistically significant (*P* <0.0001). Interpretation of the observed difference in synovial fluid spectra by PCA of the corresponding effect matrix confirmed that long-term storage results, on average, in less intense absorption peaks.

Based on the outcomes of this exploratory analysis, which indicated that sample ageing could result in small but significant differences in the spectral signature, in a second stage of the study we wanted to verify if the analysis of long-term stored samples could still provide reliable predictions and, if so, whether models built on the two set of samples (short- and long-term stored) resulted in the same set of putative markers. At first the spectra collected on the serum samples were considered. The quality of the predictive models was evaluated by considering the figures of merit calculated on the outer loop of the rDCV procedures, which correspond to the classification performances on samples not used for model building or optimization. Moreover, since rDCV uses resampling, this allows to estimate, for each figure of merit, not only a single value but also a confidence interval (in particular, in the remainder of the text, 95% confidence level will always be considered). The predictive models for discriminating serum OA samples (n =50) from controls (n=49) built on the spectra from short-term storage showed sensitivity, specificity, and accuracy of 87.3±3.7%, 88.9±2.4% and 88.1±2.3% respectively. The predictive models based on the same serum samples after long-term storage showed sensitivity, specificity and accuracy of 92.5±2.6%, 97.1±1.7% and 94.8±1.4% respectively. In both cases, the very good discriminant ability of the models can be graphically visualized in **Figure 1**, where the mean scores (and the corresponding confidence intervals) of the samples along the single canonical variate (discriminant direction) of the models are displayed. In both cases, it can be observed how almost all the OA samples have positive scores along the direction, whereas many of the controls have negative coordinates.

**Figure 1:**
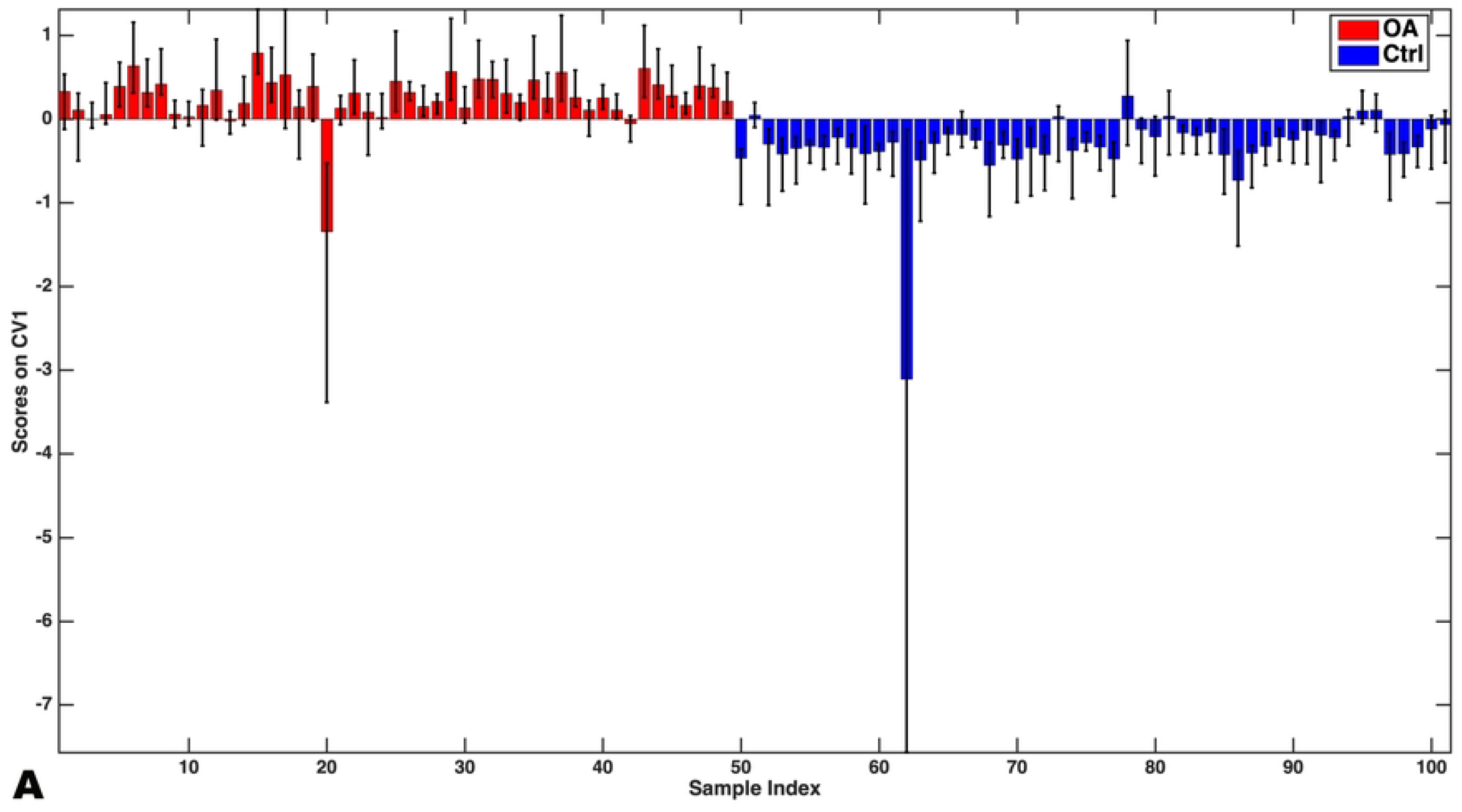
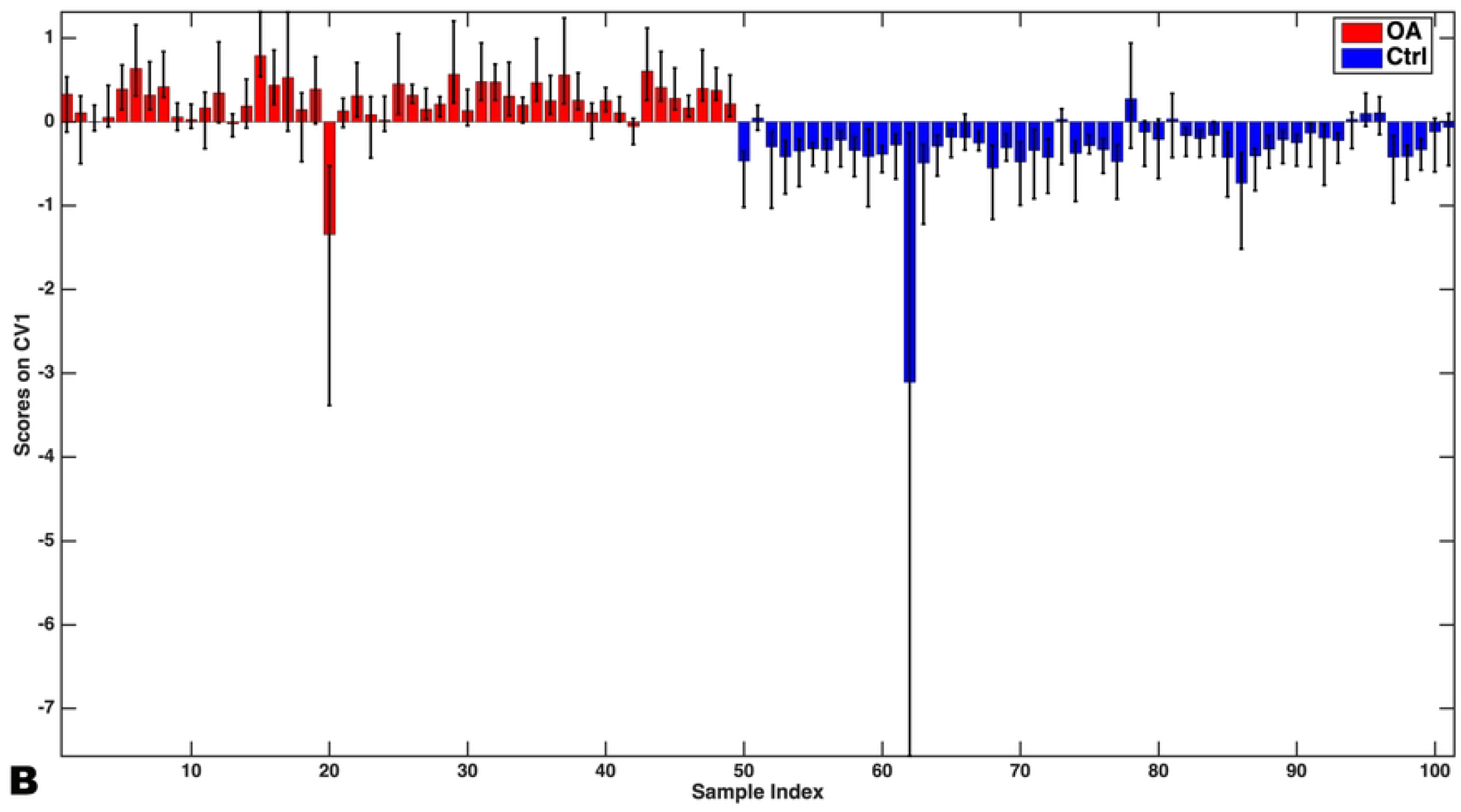
Double-cross-validated projections of the outer loop serum spectral variables onto the only canonical variate of the classification model showing the difference in the values of the scores (bars indicate mean and whiskers the corresponding 95% confidence intervals) between OA and control serum samples after short-term storage (A) and long-term storage (B). CV, canonical variate; OA, osteoarthritis; Ctrl, control.

Having verified that both models provide accurate discrimination between OA and control samples, inspection of the spectral variables mostly contributing to the observed differentiation was conducted by analyzing the values of the variable importance in projection (VIP) indices. The VIP indices summarize the contribution of each experimental variable to the PLS-DA model and are normalized so that a greater-than-one criterion can be used to assess their significance. Accordingly, the wavenumbers mostly responsible for the observed differences between OA and control serum samples at each time point are presented in **Figure 2**.

**Figure 2.**
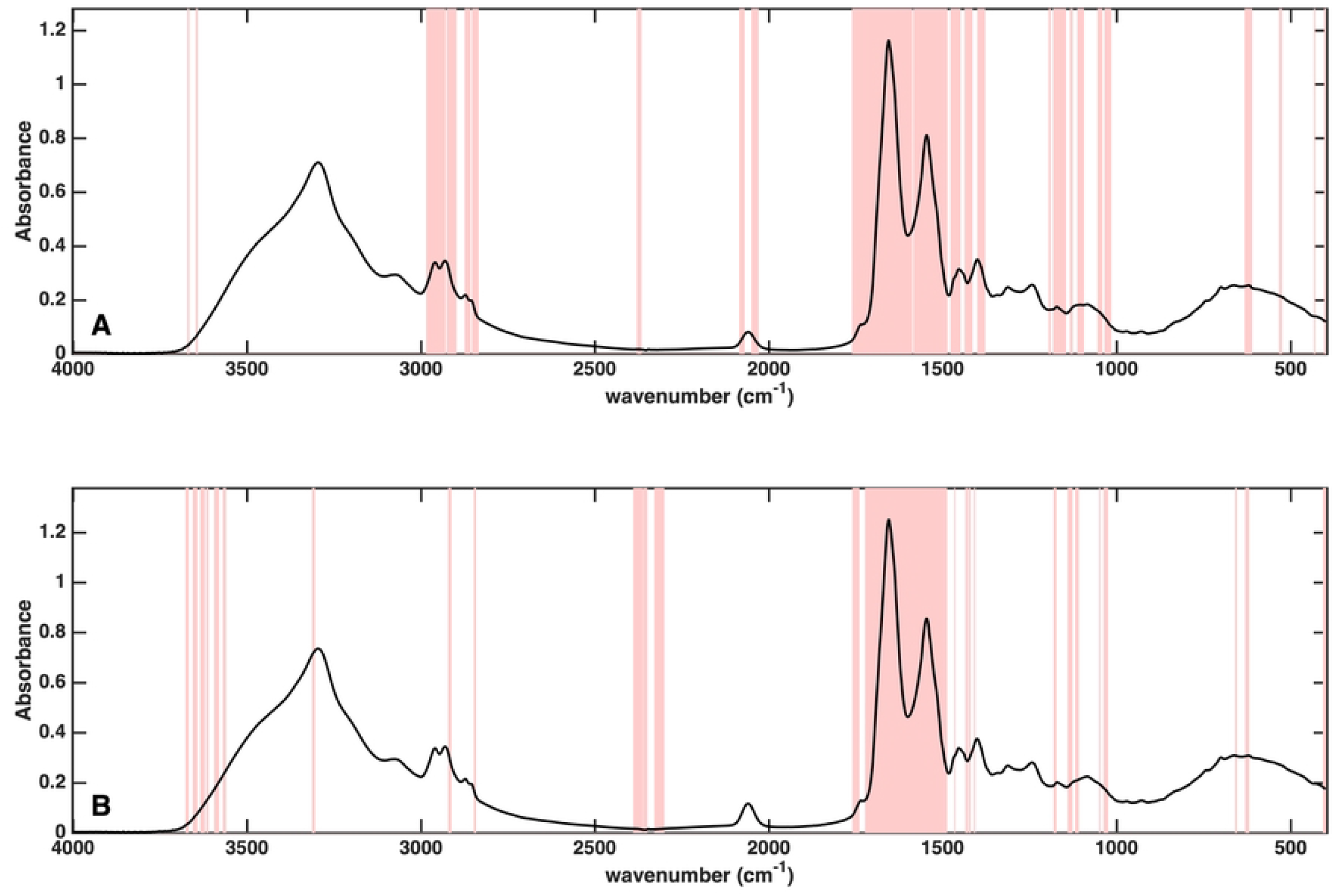
Mean spectra of serum samples from short-term (A) and long-term storage (B) with highlighted lines (pink) outlining wavenumbers responsible for the observed differences in the spectra.

The analysis was then repeated for the synovial fluid samples based on the OA (n=51) and control (n=52) groups. The classification model based on short-term storage synovial fluid spectra had sensitivity, specificity, and accuracy of 97.3±1.6%, 89.4±2.6% and 93.4±1.6% respectively, when evaluated on the outer loop of the rDCV procedure. The predictive models based on the same synovial fluid samples after long-term storage showed sensitivity, specificity, and accuracy of 95.7±2.1%, 95.7±0.8% and 95.8±1.1% respectively. Also in this case, the very good discriminant ability of the models can be graphically visualized in **Figure 3**, where the mean scores (and the corresponding confidence intervals) of the samples along the single canonical variate (discriminant direction) of the models are displayed. On the other hand, the wavenumbers responsible for the observed differences between OA and control SF samples at each time point, identified by inspection of the VIP indices calculated from the corresponding PLS-DA models, are presented in **Figure 4**.

**Figure 3:**
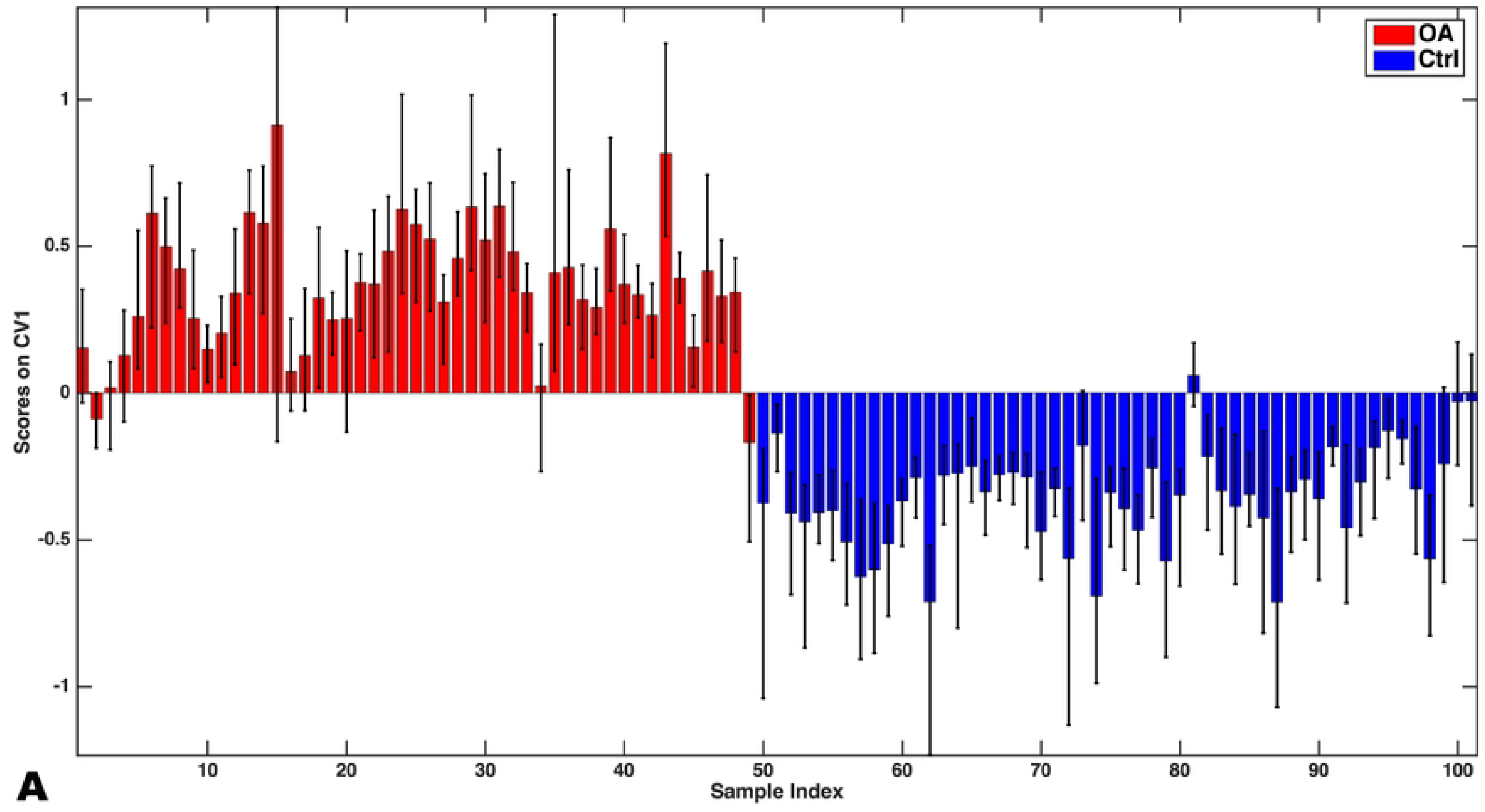
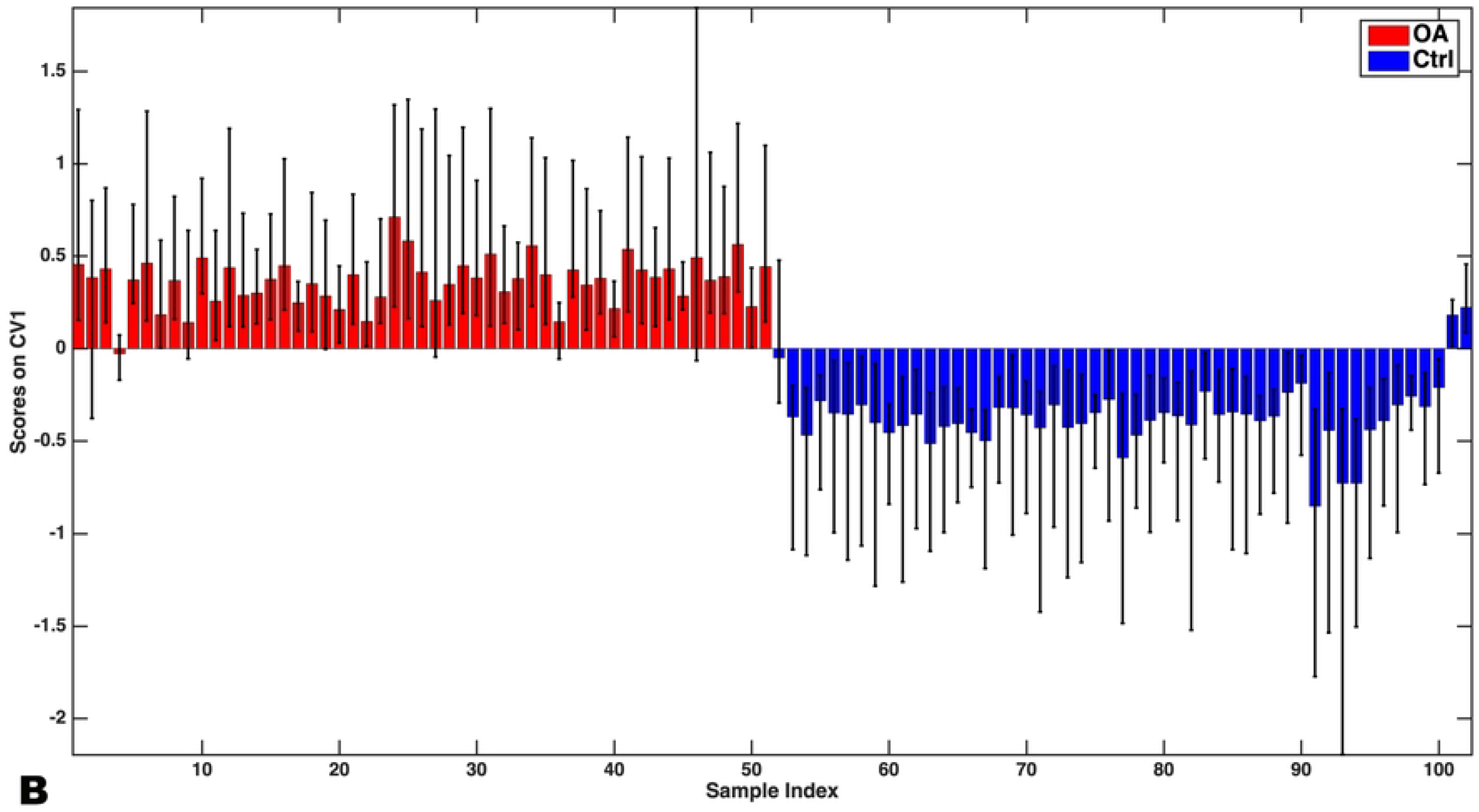
Double-cross-validated projections of the outer loop of synovial fluid spectral variables onto the only canonical variate of the classification model showing the difference in the values of the scores (bars indicate mean and whiskers the corresponding 95% confidence intervals) between OA and control synovial fluid samples after short-term storage (A) and long-term storage (B). CV, coefficient variable; OA, osteoarthritis; Ctrl, control.

**Figure 4.**
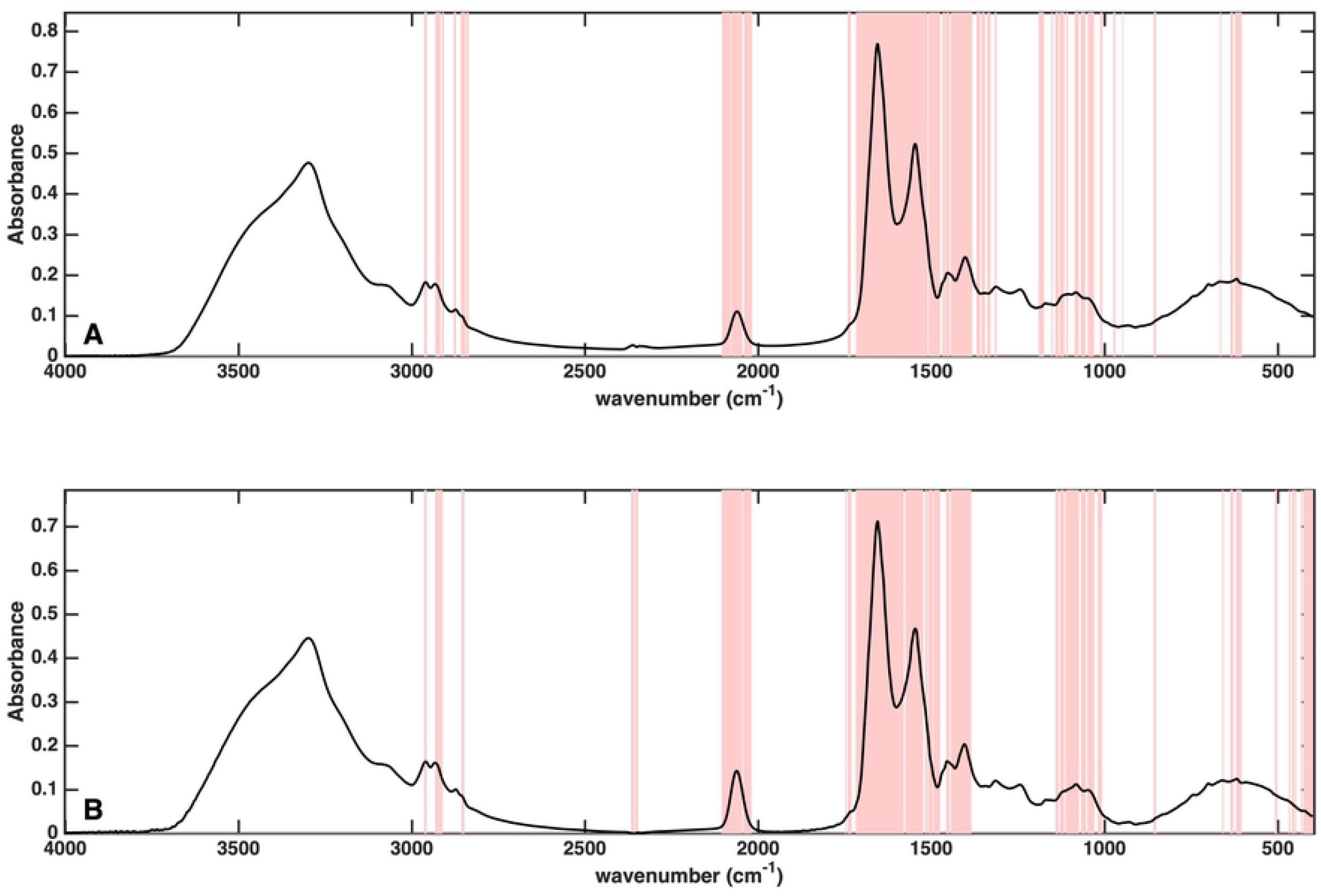
Mean spectra of synovial fluid samples from short-term (A) and long-term storage (B) with highlighted lines (pink) outlining wavenumbers responsible for the observed differences in the spectra.

## 4. Discussion

This is the first study evaluating impact of long-term sample storage on the quality of MIR spectra of serum and synovial fluid samples used for discriminating OA from control samples. Both serum and synovial fluid samples spectra had statistically significant differences in spectral variables due to aging of the samples. However, the contributions of these differences to the overall spectral variability for both serum and synovial fluid were relatively small (i.e., <3%). It is important to note that despite the statistically significant differences between the spectra of samples from short-term versus long-term storage, these differences were not a significant component of the observed differences between OA and control samples. This is reflected in the continuously high performance of the predictive models (> 87% for sensitivity, specificity and accuracy) for both serum and synovial fluid samples despite the time of measurements. Visual assessment of spectra from different time points for serum and synovial fluid samples (Figure 2 and 4) also shows how there is a high consistency in the variables identified as putative spectroscopic markers of OA between the two models, thus confirming that even if prolonged storage time can result in small but significant spectral differences, these differences do not affect the reliability and the interpretation of classification models built on the data. This finding supports use of biobanking of samples for ~7 years if spectral fingerprints are used for discriminating between OA and control samples.

This difference between the levels of variability due to aging in the samples (i.e., 1.61% in serum versus 2.98% in synovial fluid) may be partly due to the types of molecules in the synovial fluid that may be more vulnerable to freezing and degradation. But more importantly the sample storage protocols may have a role in the observed differences. Serum is cell-free whereas the synovial fluid preparation in this study involved freezing the sample without any further processing (i.e., no centrifugation or separation of cellular components pre-freezing). Therefore, presence of cellular components and their unique degradation patterns may have resulted in the greater variability between the synovial fluid spectra based on time in frozen state. The influence of different sample collection, processing, and freezing methods on results of metabolomics studies using synovial fluids have developed optimized protocols to minimize loss of data during measurements(24, 25). Therefore, if the dried-film FTIR spectroscopy technique used in this study is to be compared with metabolomics and proteomic techniques, the sample processing and storage techniques must be uniform to allow direct comparisons.

This study shows small but acknowledgeable impact of long-term storage that needs to be taken into account when utilizing this methodology for prospective clinical trials where samples are collected over years particularly for chronic and insidious disease processes such as OA. In such long prospective studies or when biobanked samples are utilized, batch analysis of the samples is preferred to reduce other variations in testing conditions (e.g., operator variation, environmental variables, etc.).Therefore, ensuring that confounding variables such as sample aging is not significantly impacting test results is paramount. The results of this study cannot be extrapolated to spectral patterns of other disease processes, as degradation of sample components unique to other disease processes may not follow the same pattern as in OA or controls. Additionally, the results of this study cannot be used to directly identify the molecular changes responsible for the observed difference in the spectral fingerprint. However, the degradation of both serum and SFs samples appears to be relatively consistent between the time points in this study, resulting in less intense absorbance peaks. The absorbance peaks of the spectra are directly related to the concentration of the sum of all molecular bonds that are MIR active at each wavenumber (i.e., Beer’s law)(13). Therefore, a decrease in peak intensity in face of unchanged testing conditions (e.g., sample dilution, spectrometer settings) is reflective of degradation of these MIR active molecular bonds. Quantitative evaluation of differences in molecules and compounds in these biological samples from different time points requires prospective studies that perform quantitative assays (e.g., proteomics, metabolomics). The identified molecules with differences over time can then be matched with the MIR peaks at wavenumbers that correspond to their molecular bonds. The main intent of the FTIR spectroscopy of dried films technique used in this study is as a cost effective, sample sparing, adjuvant free, screening tool to distinguish between samples with or without OA rather than identify the underlying causes of the observed differences(26).

Another limitation of this study is lack of additional, shorter intervals between measurements, and longer storage period to assess possible limits for sample storage time and trends as to when the sample variability due to degradation due to aging overwhelms the differences based on disease state. Future studies, can prospectively store samples and sequentially measure MIR spectra over shorter intervals (e.g. six months intervals) to document these changes. Additionally, some of the serum and synovial fluid samples in this study had undergone up to two freeze thaw cycles due to limited sample volumes available and the impact of these additional freeze thaw cycles on the overall observed changes using the FTIR spectroscopy methodology in this study cannot be separated. The impact of number of freeze thaw cycle on serum and synovial samples have been shown to have variable impact on the results based on measured variables and methods of measurement(27, 28). However, the fact that despite these confounding factors, the overall changes observed do not significantly impact the predictive modelling for discriminating OA from controls shows merit in this technique despite the limitations. Future studies, can evaluate impact of number of freeze/thawing cycles on the quality of MIR spectral patterns obtained using the methodology in this study.

## 5. Conclusions

In conclusion, storing serum and synovial fluid samples of dogs with knee OA and controls in −80°C results in changes in the spectral patterns after ~5 years. However, these changes have minimal impact on the ability to use the spectral fingerprints of these samples after long-term storage for discriminating between OA and control samples. Future studies can evaluated measured MIR spectra of a samples in storage prospectively at shorter intervals to establish trends of sample degradation over longer follow up times and set maximal limits on sample viability for biobanking purposes.

## Abbreviations

OA: osteoarthritis
FTIR spectroscopy: Fourier-transform infrared spectroscopy
MIR: Mid-infrared
KSCN: potassium thiocyanate
MSCA: Multilevel simultaneous component analysis
rDCV: Repeated double cross-validation
PLS-DA: Partial least squares discriminant analysis
VIP: Variable importance in projection

## Acknowledgements

We would like to thank Ms. Cynthia Mitchel for her assistance in performing the spectral data acquisition at the University of Prince Edward Island.

## CRediT Author statement

**Sarah Malek:** Conceptualization, Methodology, Data curation, Investigation, Formal analysis of non-spectral data, Writing-original draft preparation, project administration, Funding acquisition (personal faculty start up fund at institution) **Federico Marini:** Formal analysis of spectral data, Writing-review and editing, **J. Trent McClure:** Resources, data acquisition, Writing-review and editing.

